# Particle Biology: A Perspective on a First-Principles Theory of Life

**DOI:** 10.64898/2026.05.17.725705

**Authors:** Peng Wang, Wen Li, Yishuang Cui, Hongjiao Wu, Junqing Gan, Weinan Yao, Ye Jin, Yanna Bi, Yanlei Ge, Guogui Sun

**Author notes:** Correspondence: Yanlei Ge Guogui Sun. These authors contributed equally to this work (Peng Wang, Wen Li).

## Abstract

This Perspective formally proposes Particle Biology as a unifying theoretical framework to address the critical bottleneck in current life science research. Current life science research has reached a critical bottleneck. While the field has advanced to the study of 3D genomic spatial configurations and chromosomal organization, it remains largely descriptive and confined to the macromolecular level. This approach lacks a first-principles understanding of the underlying physical forces that drive biological processes. This Perspective formally proposes Particle Biology as a unifying theoretical framework. We establish an axiomatic system positing that life phenomena are fundamentally emergent spatiotemporal patterns of electromagnetic forces among atoms, electrons, and nuclei operating far from thermodynamic equilibrium. By defining biological states through the Biological Hamiltonian and mapping biochemical pathways to multidimensional Potential Energy Surfaces (PES), we bridge the gap between descriptive biology and predictive physics. We categorize core research technologies into three modalities—seeing, computing, and controlling particles—facilitated by advancements in Cryo-EM, AlphaFold 3, and Boron Neutron Capture Therapy (BNCT). Ultimately, the trajectory of molecular biology has evolved from cells to DNA and onto the 3D spatial genome, yet it cannot go deeper within current paradigms. The next logical evolution is to move beyond the macromolecular bottleneck to focus on the electromagnetic interactions between atoms and ions—the true ‘Particle Biology’ level—to redefine disease and intervention.

## 1. Introduction: The Bottleneck of Descriptive Biology

First-principles thinking requires deriving complex phenomena from the most fundamental axioms. Erwin Schrödinger pioneered this approach in the life sciences, conceptualizing the cell through the lens of physics[1]. Today, molecular biology has achieved remarkable success in mapping the flow of genetic information and complex signaling cascades[2]. However, this success has led the field to a fundamental bottleneck: it remains largely a descriptive science. We know what molecules exist and where information flows, but we lack a bottom-up, predictive understanding of how fundamental physical forces orchestrate these molecular events.

The essence of chemical reactions within the human body is the continuous rearrangement of electron clouds — the precise operation of electromagnetic forces governed by quantum mechanics[3]. This Perspective introduces Particle Biology, a framework asserting that the essence of biological research must shift toward the fundamental interactions between particles. By stripping away biological complexity to reveal the underlying quantum and thermodynamic logic, we aim to establish a unified perspective that guides computational prediction, experimental design, and the next generation of therapeutic interventions[4-8].

## 2. The Axiomatic Foundations of Particle Biology

To transition from a descriptive discipline to a predictive physical science, Particle Biology establishes three foundational axioms that govern all cellular processes.

### 2.1 Axiom I: Material Identity and Electromagnetic Dominance

Living organisms possess no unique “vital force.” The constituents of life (quarks, electrons, atomic nuclei) strictly obey the Standard Model of particle physics. Within the biological temperature and energy scales, the electromagnetic force is the absolute dominant interaction dictating biomolecular structure, recognition, and transformation[2, 9, 10].

### 2.2 Axiom II: Thermodynamic Dissipation

Life is an open system operating far from thermodynamic equilibrium[11]. It continuously intakes low-entropy substrates to sustain local order (negentropy) while dissipating high-entropy waste into the chaotic background of Brownian motion. At the microscopic level, every biochemical pathway is strictly constrained by the Gibbs free energy equation[12]: ΔG = ΔH - TΔS

### 2.3 Axiom III: Quantum Dominance in Core Processes

Classical mechanics is insufficient to describe life’s efficiency. In critical biological events— such as enzyme catalysis, energy transduction, and mutagenesis—quantum mechanical effects (e.g., quantum tunneling, superposition, and coherence) are not merely environmental noise but the core driving mechanisms honed by evolution[13-15].

## 3. Mathematical and Physical Formulation

Under the Particle Biology paradigm, a biological complex (e.g., a receptor-ligand complex or a replisome) is not a geometric puzzle but a complex quantum many-body system.

### 3.1 The Biological Hamiltonian

The state of any biological system can be fundamentally described by its total Hamiltonian:

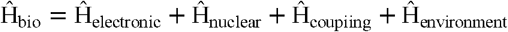

Biological function emerges from the interplay of electronic states (Ĥ _electronic_), nuclear configurations (Ĥ_nuclear_), their vibronic coupling (Ĥ_coupling_), and the stochastic thermal fluctuations of the aqueous environment (Ĥ_environment_)[16, 17].

### 3.2 The PES and Disease Definition

Biochemical reactions are trajectories across a multidimensional PES defined by the Hamiltonian. Normal physiological functions represent the system finding the globally optimal thermodynamic pathways. Consequently, disease is fundamentally a physical state: it occurs when a biological system becomes trapped in a pathological local minimum (a metastable state) on the PES, or when environmental parameters (e.g., local pH, temperature, oxidative stress) alter the electromagnetic charge distribution, thereby distorting the energy landscape itself[18, 19].

### 3.3 Beyond Extreme Reductionism: The Role of Emergence

A common critique of first-principles thinking in biology is the risk of extreme reductionism —the assumption that complex traits can be understood merely by summing the properties of isolated particles. Particle Biology explicitly rejects this. While electromagnetic forces and quantum mechanics are the fundamental drivers, life is an open, non-linear complex system. The concept of emergence is central to our framework[20]. At each hierarchical scale, the topological constraints of the interacting particles generate novel macroscopic properties that do not exist at the lower levels. For instance, the quantum entanglement of electrons dictates covalent bonding, but the network topology of these bonds (e.g., in a folded antibody) gives rise to emergent phenomena like immune memory and allosteric regulation[21]. Therefore, Particle Biology does not reduce consciousness or immunity to a single particle’s state; rather, it provides the precise physical grammar (forces and energy landscapes) through which these highly complex, emergent biological narratives are written **(Table 1**).

**Table 1.** Hierarchical Emergence from Subatomic Particles to Cellular Systems.

| Hierarchical Level | Interacting Entities | Dominant Physical/Quantum Concept | Emergent Biological Function |
| --- | --- | --- | --- |
| Subatomic[22] | Electrons, Protons | Quantum Tunneling, Superposition | Enzyme catalysis rate acceleration, directional energy capture. |
| Atomic/Molecular[23] | Atoms, Functional Groups | Electron-Phonon Coupling (He-ph) | Protein folding stability, allosteric conformational changes. |
| Macromolecular[24] | Proteins, DNA strands | Free Energy Landscapes ($\Delta G$) | Lock-and-key recognition, genetic transcription fidelity. |
| Cellular/System[25] | Organelles, Cell Networks | Non-equilibrium Thermodynamics | Immune memory, action potential propagation, cellular differentiation. |

## 4. Case Study I: Quantum Biology in Natural Systems

The explanatory power of Particle Biology is most evident where classical biochemistry fails to explain the efficiency of life.

### 4.1 Quantum Tunneling in Enzyme Catalysis

Classical transition state theory cannot fully account for the extraordinary rate accelerations achieved by enzymes like alcohol dehydrogenase. The transfer of hydrogen is not a classical journey over the activation energy barrier, but involves quantum tunneling—where the wave-like nature of the proton or electron allows it to pass through the potential barrier[15, 21]. This is experimentally validated by anomalous kinetic isotope effects when substituting hydrogen with deuterium, proving that life harnesses subatomic quantum probability to achieve catalytic perfection[26].

### 4.2 Quantum Coherence in Photosynthesis

The capture and transfer of solar energy in the Fenna-Matthews-Olson (FMO) complex occurs with near-100% efficiency. Excitons do not randomly hop between bacteriochlorophyll molecules; rather, they exist in a quantum superposition state, simultaneously exploring multiple energy pathways via quantum coherence to find the thermodynamically optimal route to the reaction center[27, 28].

## 5. Case Study II: Smart Hydrogels as Engineered Particle Systems

If natural biology is the evolutionary optimization of particle interactions, smart hydrogels represent the human engineering of these precise principles, validating the reductionist perspective of Particle Biology.

### 5.1 Programming Orbital Interactions

The “intelligence” of a tumor-microenvironment responsive hydrogel lies in chemical bonds acting as molecular switches. The cleavage of disulfide bonds by glutathione is a nucleophilic substitution governed by FMO theory. The lone pair electrons of the thiolate anion precisely attack the σ ∗ antibonding orbital of the disulfide bond. By engineering theΔG of this reaction, we ensure macroscopic material degradation is strictly triggered by microscopic orbital overlap[29, 30].

### 5.2 Macroscopic Control via External Energy Fields

Advanced hydrogels integrate superparamagnetic nanoparticles. An external alternating magnetic field forces the electron spins to continuously align and relax, dissipating localized heat (hyperthermia). This thermal energy increases local Brownian motion, overcoming the activation energy barrier for polymer dissociation. This engineered cascade—from macroscopic magnetic fields to nanoscale spin dynamics to molecular bond cleavage—perfectly illustrates the top-down control of the biological Hamiltonian[31, 32].

## 6. Falsifiable Predictions

A valid scientific theory must be able to predict the unknown. Particle Biology moves beyond descriptive models to offer concrete, falsifiable predictions:

### 6.1 Ab Initio Binding Energy Prediction

Without relying on empirical crystal structure databases, Particle Biology posits that utilizing density functional theory (DFT) and quantum-mechanics/molecular-mechanics (QM/MM) calculations can precisely predict the binding free energy (ΔG) of de novo designed drug molecules to target proteins, predicting efficacy solely from electron density distributions[33, 34].

### 6.2 Quantitative Quantum Kinetics

For specific hydrogen-bond networks in mutated enzymes, the theory predicts that the loss of catalytic efficiency is not due to gross structural mismatch, but a calculable reduction in the quantum tunneling probability, which can be quantitatively verified via low-temperature KIE experiments[35, 36].

### 6.3 Executable Computational Paths: From QM/MM to AI Force Fields

To transition from theory to actionable science, Particle Biology relies on specific, multi-scale computational pipelines. Historically, modeling an entire protein using pure quantum mechanics was computationally prohibitive. Today, QM/MM bridges this gap by treating the active site (where electron rearrangement occurs) with high-level QM, while simulating the surrounding protein scaffold using classical MM force fields. Furthermore, the advent of Neural Network Potentials (NNPs), such as DeepMD, represents a paradigm shift. By training AI models on ab initio DFT data, NNPs achieve quantum-level accuracy at the speed of classical molecular dynamics[37-40]. This computational triad (QM/MM, DFT, NNPs) makes the calculation of multi-dimensional energy landscapes practically executable, allowing researchers to predict reaction rates and binding affinities strictly from particle-level interactions (**Table 2**).

**Table 2.** Multiscale Computational Pipeline: From Quantum Accuracy to Macroscopic Feasibility.

| Computational Method | Resolution Limit | Primary Application in Particle Biology | Key Advantage |
| --- | --- | --- | --- |
| DFT [41] | ~Hundreds of atoms | Ab initio calculation of electron densities and bond cleavage. | High accuracy for fundamental quantum states; no empirical parameters. |
| QM/MM Hybrid Models[42] | ~100,000+ atoms | Modeling localized enzyme active sites within a full protein scaffold. | Balances quantum accuracy with macroscopic computational feasibility. |
| NNPs [40, 43] | Millions of atoms | Simulating massive biological complexes with quantum-level accuracy. | AI-driven; achieves ab initio accuracy at molecular dynamics speeds. |
| Coarse-Grained MD[40] | Entire cell regions | Simulating long-timescale Brownian motion and complex self-assembly. | Maps broad Free Energy Landscapes; bridges micro to macro behavior. |

## 7. Technology Classification and Clinical Intervention

Under this paradigm, all cutting-edge technologies converge into three fundamental modalities (**Table 3**).

**Table 3.**
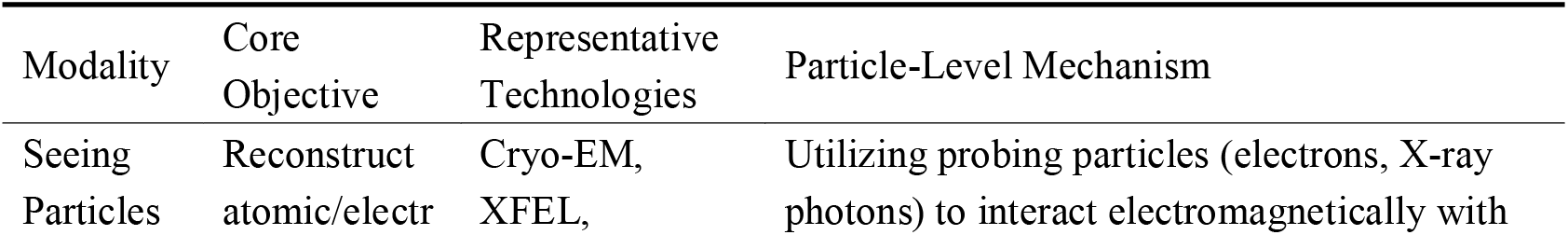
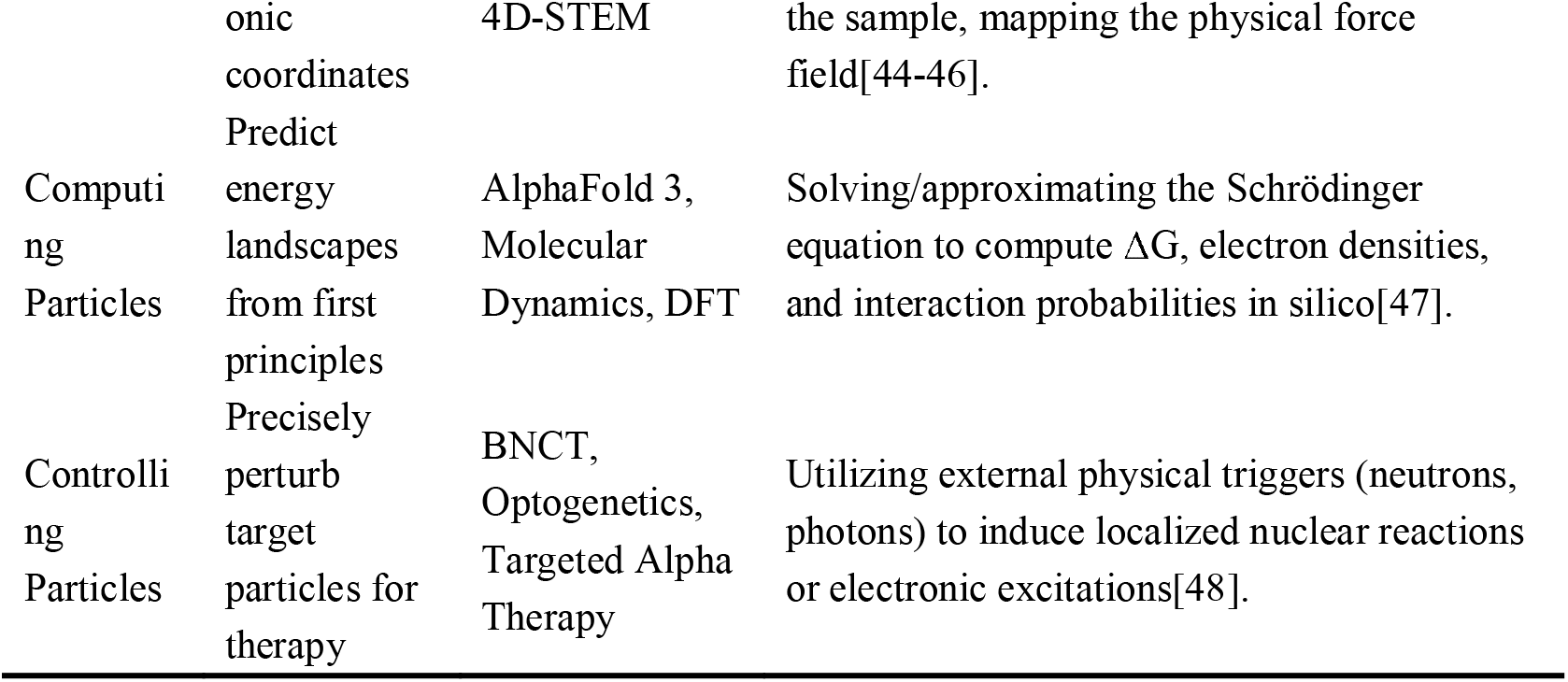
The Three Modalities of Particle Biology: From Seeing to Controlling.

### 7.1 The Future of Intervention: Particle Therapy

Traditional pharmaceuticals modulate downstream molecular cascades. Particle Biology advocates for intervening at the foundational physics level. BNCT exemplifies this: thermal neutrons trigger localized nuclear fission ^10^B+n→^7^Li+α exclusively within cancer cells[49, 50]. It bypasses biological resistance (pathway mutations) entirely by utilizing subatomic physics to destroy the target’s physical structure from within[51].

## 8. Conclusion: The Unfinished Revolution

From the perspective of first principles, Particle Biology strips away the illusion of biological exceptionalism, revealing life as the most elegant manifestation of electromagnetic forces operating far from thermodynamic equilibrium. It does not replace molecular biology but serves as its necessary evolution. By defining biological states through Hamiltonians and potential energy surfaces, we empower ourselves to move from merely observing biology to computing and engineering it.

The history of molecular biology has moved from cells to DNA, and now to the 3D spatial configuration of chromosomes. However, even at this peak, the current paradigm has reached a bottleneck. While current research into 3D genome spatial configurations is exquisite, the fluctuations in ‘spatial positioning’ are essentially driven by changes in electrostatic potential energy; without descending to the particle level, we remain observers of outcomes rather than controllers of causality. The next logical evolution must occur at the particle-to-particle level, particularly in cancer treatment. Future breakthroughs will not rely on modulating downstream molecular cascades but on controlling the fundamental interactions between particles. Technologies like Boron Neutron Capture Therapy BNCT already exemplify this, using subatomic physics to destroy malignant structures with absolute precision. The convergence of AI models and quantum therapies proves that the next scientific revolution has begun: we are mastering the modalities of seeing, computing, and controlling the fundamental particles of life. In doing so, we finally fulfill the legacy of Erwin Schrödinger: if life is defined by its ability to extract “negentropy” to sustain order against decay, Particle Biology provides the first precise operational manual for negentropy extraction at the quantum level.

## Declarations

### Ethics approval and consent to participate

Not applicable.

### Consent for publication

Not applicable.

### Availability of data and materials

All study data are included in the article. No other scripts or software were used. The analyzed datasets generated during the present study are available from the corresponding author upon reasonable request.

### Competing interests

The authors declare that they have no competing interests.

### Funding

This work was supported by the Natural Science Foundation of Hebei Province (Grant No.H2023209083); National Natural Science Foundation of China (Grant No.82172658); National Natural Science Foundation of China (Grant No.82472636); Tangshan Municipal Science and Technology Plan Project (Grant No.24150218C); Government Funded Clinical Medicine Outstanding Talent Training Program (Grant No.ZF2024193); Tangshan Municipal Science and Technology Plan Project (Grant No.25150220E); Special Project of Joint Innovation for Medicine, Research and Enterprise in Hebei Province (Grant No.LH20250016); Scientific Research Project of Universities in Hebei Province (Grant No.CXZX2025020).

### Authors’ contributions

Guogui Sun and Yanlei Ge: Conceptualization. Wen Li, Peng Wang, Hongjiao Wu and Yishuang Cui: Writing-Original draft preparation. Junqing Gan, Weinan Yao and Ye Jin: Writing - Review & Editing., Guogui Sun: Funding acquisition. Yanlei Ge and Yanna Bi: Supervision.

## Acknowledgements

Not applicable.

---

**Figure 1.**
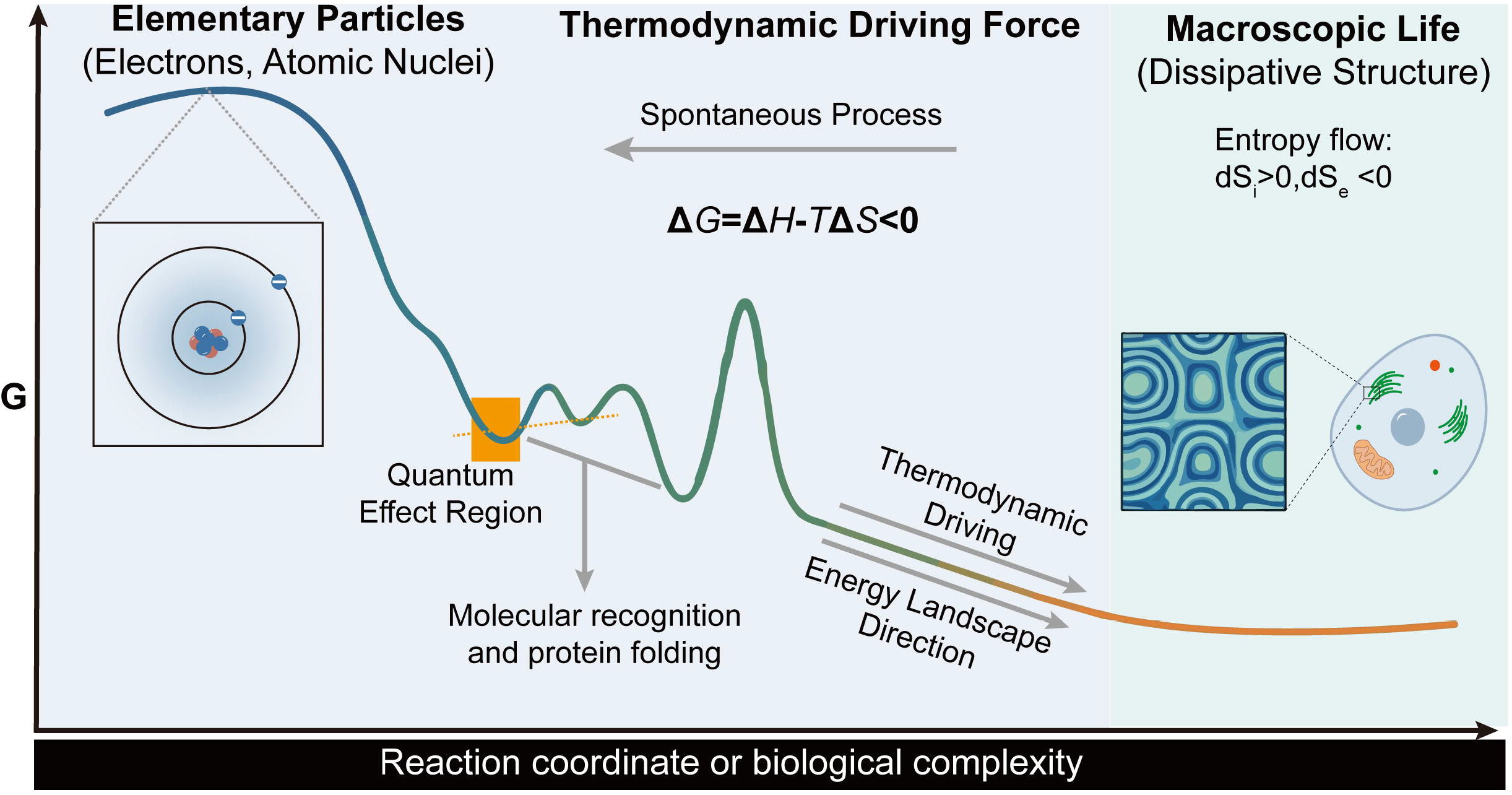
Energy Landscape from Elementary Particles to Macroscopic Life. The schematic illustrates how biological processes are fundamentally trajectories across a multidimensional energy landscape governed by the Gibbs free energy equation (ΔG = ΔH − TΔS < 0). At the microscopic scale, elementary particles (electrons and atomic nuclei) interact via electromagnetic forces to form the quantum many-body system of biomolecules. The “Quantum Effect Region” highlights where quantum mechanical phenomena—such as quantum tunneling and coherence—govern molecular recognition and protein folding. As complexity increases along the reaction coordinate, the system navigates through local energy minima toward the global thermodynamic optimum. The right panel depicts the emergence of macroscopic life as a dissipative structure, characterized by negative entropy exchange with the environment (dS□ > 0, dS□ < 0), consistent with the thermodynamic dissipation axiom of Particle Biology.

**Figure 2.**
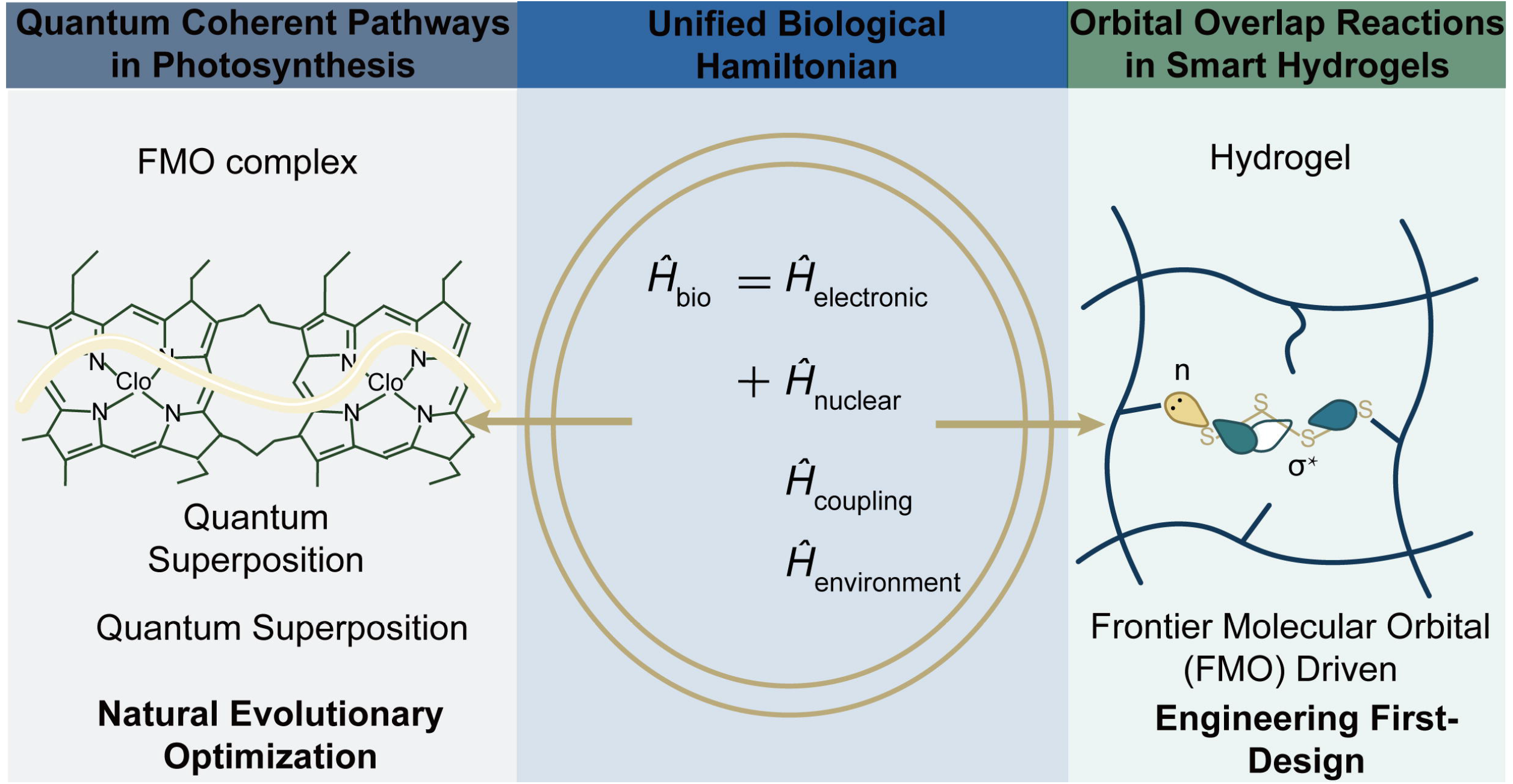
Unity of Natural and Artificial Systems (Unified Bio-Hamiltonian). The central equation, Ĥ_bio = Ĥ_electronic + Ĥ_nuclear + Ĥ_coupling + Ĥ_environment, defines the total Hamiltonian for any biological system within the Particle Biology framework. Left pane: In the Fenna-Matthews-Olson FMO photosynthetic complex, excitons exist in quantum superposition states, simultaneously exploring multiple energy transfer pathways via quantum coherence to achieve near-100% efficient energy transfer—representing natural evolutionary optimization of particle interactions. Right panel: In tumor-microenvironment-responsive smart hydrogels, the cleavage of disulfide bonds by glutathione is driven by FMO overlap, where the thiolate lone pair attacks the σ* antibonding orbital. This exemplifies human-engineered control of particle interactions through first-principles design. Together, these natural and artificial systems validate the unifying explanatory power of the Biological Hamiltonian.

**Figure 3.**
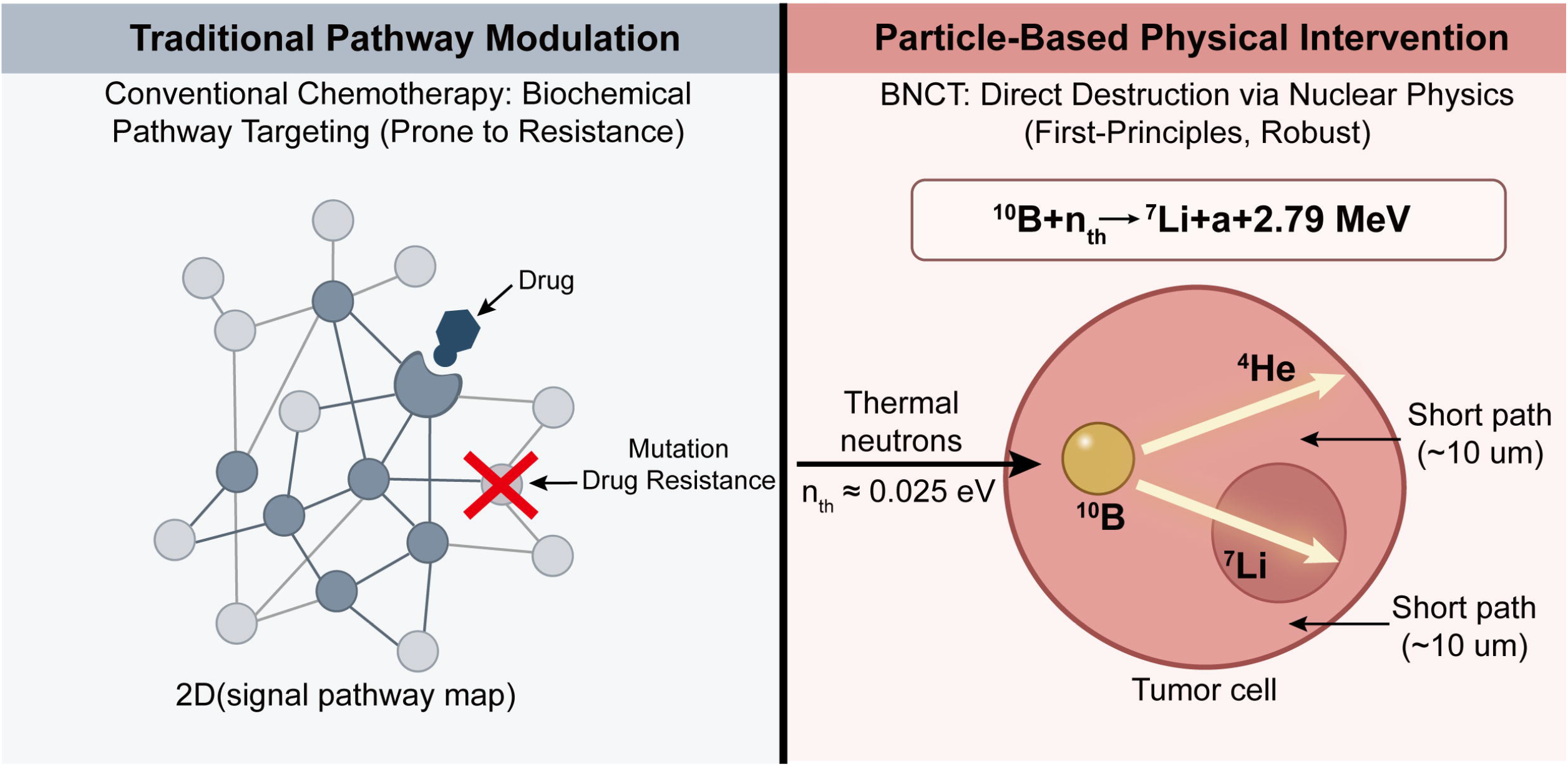
Particle Intervention Schematic (BNCT as an Example) - Physical Precision Strike vs Traditional Pathway Modulation. Left panel: Conventional chemotherapy operates at the descriptive molecular biology level, targeting nodes within two-dimensional signal pathway maps. This approach is inherently prone to drug resistance, as pathway mutations (red cross) can bypass the drug-target interaction. Right panel: In contrast, Particle Biology advocates for first-principles intervention at the physical level. BNCT exploits the nuclear capture reaction ^1^ □B + n□□ → □Li + α + 2.79 MeV, where thermal neutrons (n□□ ≈ 0.025 eV) trigger localized nuclear fission exclusively within tumor cells preloaded with ^1^ □B. The resulting α particles and □Li nuclei travel only ∼10 μm, delivering lethal, subcellular-precision damage that completely bypasses biological resistance mechanisms. This represents the “Controlling Particles” modality—intervening at the foundational physics level rather than modulating downstream molecular cascades.

## Notes

### Competing Interest Statement

The authors have declared no competing interest.

